# Correction of transposase sequence bias in ATAC-seq data with rule ensemble modeling

**DOI:** 10.1101/2022.12.08.519600

**Authors:** Jacob B. Wolpe, André L. Martins, Michael J. Guertin

## Abstract

Chromatin accessibility assays have revolutionized the field of transcription regulation by providing single-nucleotide resolution measurements of regulatory features such as promoters and transcription factor binding sites. ATAC-seq directly measures how well the Tn5 transpose accesses chromatinized DNA. Tn5 has a complex sequence bias that is not effectively scaled with traditional bias-correction methods. We model this complex bias using a rule ensemble machine learning approach that integrates information from many input k-mers proximal to the ATAC sequence reads. We effectively characterize and correct single-nucleotide sequence biases and regional sequence biases of the Tn5 enzyme. Correction of enzymatic sequence bias is an important step in interpreting chromatin accessibility assays that aim to infer transcription factor binding and regulatory activity of elements in the genome.

## Introduction

Chromatin accessibility assays measure the relative frequency that exogenous enzymes access DNA. Chromatin accessibility is not a direct and measure molecular features of chromatin such as transcription factor occupancy or histone modification status. However, accessibility is considered a proxy measurement of regulatory element activity irrespective of the constellation of factors bound to the DNA (Wu et al. 1979a,b). Accessible regions are enriched for transcription factor binding and histone modifications that are hallmarks of functional *cis*-regulatory elements (Boyle et al. 2008; Guertin et al. 2012; Moore et al. 2020; Tewari et al. 2012; Thurman et al. 2012). Deconvolution of genomewide chromatin hypersensitivity data to infer individual transcription factor binding events remains a challenge (Li et al. 2019a).

ATAC-seq revolutionized the chromatin accessibility field by providing a straightforward method that requires fewer than 5000 cells (Buenrostro et al. 2015). ATAC-seq leverages a hyperactive Tn5 transposase that directly inserts high throughput sequencing adapters into accessible DNA to create sequencing libraries. The analysis of ATAC-seq data requires additional considerations beyond traditional hypersensitivity assays because the molecular biology of transposase function is distinct compared to DNase and other enzymes that are used to measure chromatin accessibility (Buenrostro et al. 2013; Smith et al. 2021).

Although using Tn5 to measure chromatin accessibility is experimentally easier than using DNase, the dimeric Tn5 enzyme recognizes a wider region when interacting with DNA and this is reflected in the sequence bias of Tn5 (Li et al. 2019b). Characterizing and correcting enzymatic sequence bias is an essential step for accurate interpretation of chromatin accessibility data (He et al. 2014; Koohy et al. 2013; Sung et al. 2014). Strategies to correct biases use a variety of sequence inputs to build models, including k-mer instances, position weight matrices, and long stretches of DNA. DNase bias can be directly scaled based on the 6-mer sequence centered on the DNase cleavage site (Martins et al. 2018; Schwessinger et al. 2017; Wang et al. 2017; Yardımcı et al. 2014). The advantage of direct k-mer scaling is the simplicity: reads are scaled by the expected / observed k-mer count ratio. If a k-mer is found less often than expected by chance, then the read is scaled to a higher value. However, direct k-mer scaling does not effectively correct Tn5 biases from ATAC-seq data (Karabacak Calviello et al. 2019; Martins et al. 2018; Schwessinger et al. 2017), so other methods were developed to correct Tn5 bias. `ATACorrect` scales individual ATAC-seq reads based upon a dinucleotide weight matrix representing Tn5 bias (Bentsen et al. 2020). Similar to weight matrix scaling, `HINT-ATAC` employs position dependency models, which account for interactions between weight matrix positions to model ATAC-seq bias (Li et al. 2019b). `SELMA` encodes k-mer data into a simplex vector, which is incorporated into a Hadamard matrix for all mono and dinucleotide combinations. These data are input into into a linear regression model to capture and correct both DNase and Tn5 bias (Hu et al. 2022). More sophisticated methods that correct ATAC-seq data are often less interpretable than k-mer and weight matrix scaling. `Seqbias` trains a Bayesian network to encode nucleotide preferences using a 40 base window centered on the Tn5 recognition site (Viswanadham et al. 2019). `MsCentipede` uses naked DNA cleavage to train a Bayesian multi-scale model of a Poisson distribution of reads to account for sequence bias (Raj et al. 2015). Several methods employ convolutional neural networks to account for intrinsic sequence biases (Ansari et al. 2020; Yang and Henao 2022).

We previously found that direct k-mer scaling corrects the majority of nuclease sequence bias. We developed `seqOutBias` to implement k-mer scaling for molecular genomics data. The `seqOutBias` package is stand-alone software that specializes in sequence bias correction for a range of k-mer lengths and gapped k-mers. `SeqOutBias` output files can be piped into programs that specialize in peak calling and footprint inference (Gaspar 2018), but `seqOutBias` performs poorly with ATAC-seq data. Here, we expanded the `seqOutBias` package to accommodate ATAC-seq data by coupling `seqOutBias` output to a rule ensemble modeling framework that effectively scales individual ATAC-seq reads to correct Tn5 bias. This modeling approach captures complex interactions between k-mers and quantifies the importance of individual positions that contribute to overall Tn5 bias. Moreover, the importance of k-mers and positions are intrepretable locally for each position in the genome, because the rules and terms applied to each aligned read are explicit. This reproducible workflow efficiently corrects single-nucleotide resolution Tn5 sequence bias and addresses regional baseline bias determined by GC content. To facilitate bias correction of chromatin accessibility we developed a workflow that can be applied to existing high-throughput sequencing analysis pipelines: https://github.com/guertinlab/Tn5bias.

## Results

### Tn5 sequence bias is more extensive than other nuclease biases

ATAC-seq measures how well Tn5 transposase accesses DNA in chromatin, which is a proxy measurement for regulatory element activity. Although chromatin condensation is the main contributor that influences Tn5-mediated DNA cleavage, the local sequence content biases transposition activity (Buenrostro et al. 2013). A dimeric Tn5 complex nicks opposite strands of DNA 9 base pairs apart, and each monomer inserts a sequencing-compatible adapter at each nick position (Figure S1A) (Reznikoff 2003, 2008). The interaction of Tn5 with a precise genomic region results in reads aligning to two converging genomic coordinates 9 bases apart on opposite strands. Therefore the field shifts the reads aligning to either reference genome strand so that a common position anchors the Tn5 recognition site regardless of the orientation of the adapter. We refer to the center nucleotide of this 9 base insertion site as the **central** Tn5 recognition **base**, in contrast to a traditional nucleases which cleavage a single position between two adjacent bases. Shifting ATAC-seq reads results in base-pair resolution data that precisely identifies the center of the Tn5 recognition site.

Detection of a single Tn5-inserted adapter necessitates an independent Tn5 insertion within approximately 500 bases. Only half of the Tn5 insertion events are capable of amplification, even when an independent insertion occurs within 500 bases (Figure S1B). To specify this central base, we shift the forward-strand aligned reads downstream by 4 bases and we shift the reverse-strand aligned reads upstream by 4 bases. Most current ATAC-seq analysis workflows shift the reverse-strand aligned reads upstream by 5 bases so that the data looks more similar to the chromatin accessibility predecessor DNase-seq. However, this approach requires an independent shift of one base for reverse-aligned sequences to ensure that a single DNA cleavage event is specified by a single genomic coordinate (Vierstra and Guertin 2021).

All enzymes that recognize nucleic acids as substrates have sequence biases. These sequence biases are quantified as the observed frequency of each base at individual positions relative to a cleavage site (or the centrally recognized base in the case of transposases) compared to the expectation of random genomic cleavage. Enzyme bias motifs are represented by a position probability matrix that reports the fraction of each base observed at each position relative to the cut site, which is most easily visualized as a seqLogo (Gavin E. Crooks and Brenner 2004; Schneider and Stephens 1990). We confirm that Tn5 favors CG-rich DNA and the seqLogo is a reverse complement palindrome, which supports Tn5 recognizing DNA as a dimer (Figure 1A) (Buenrostro et al. 2013; Viswanadham et al. 2019). The nucleases Benzonase, Cyanase, DNase, and MNase have distinct biases that span fewer positions compared to Tn5 (Figure 1A) (Grøntved et al. 2012; Iwata-Otsubo et al. 2017; Lazarovici et al. 2013). In addition to Tn5 having a wider bias window, the cumulative information content within the window is highest for Tn5 compared to the nucleases (Figure 1A). Importantly, we consider background nucleotide frequency when calculating information content. If we did not background-correct, then random cleavage would be interpreted as an AT bias because A/T bases account for 59% of the genome (Figure S2A). Previous methods that accurately correct enzyme sequence biases use k-mer scaling (Martins et al. 2018). However, k-mer scaling does not effectively correct Tn5 bias because of the wide bias window and the limitation that large k-mers occur rarely in both the observed and expected k-mer counts.

**Fig. 1.**
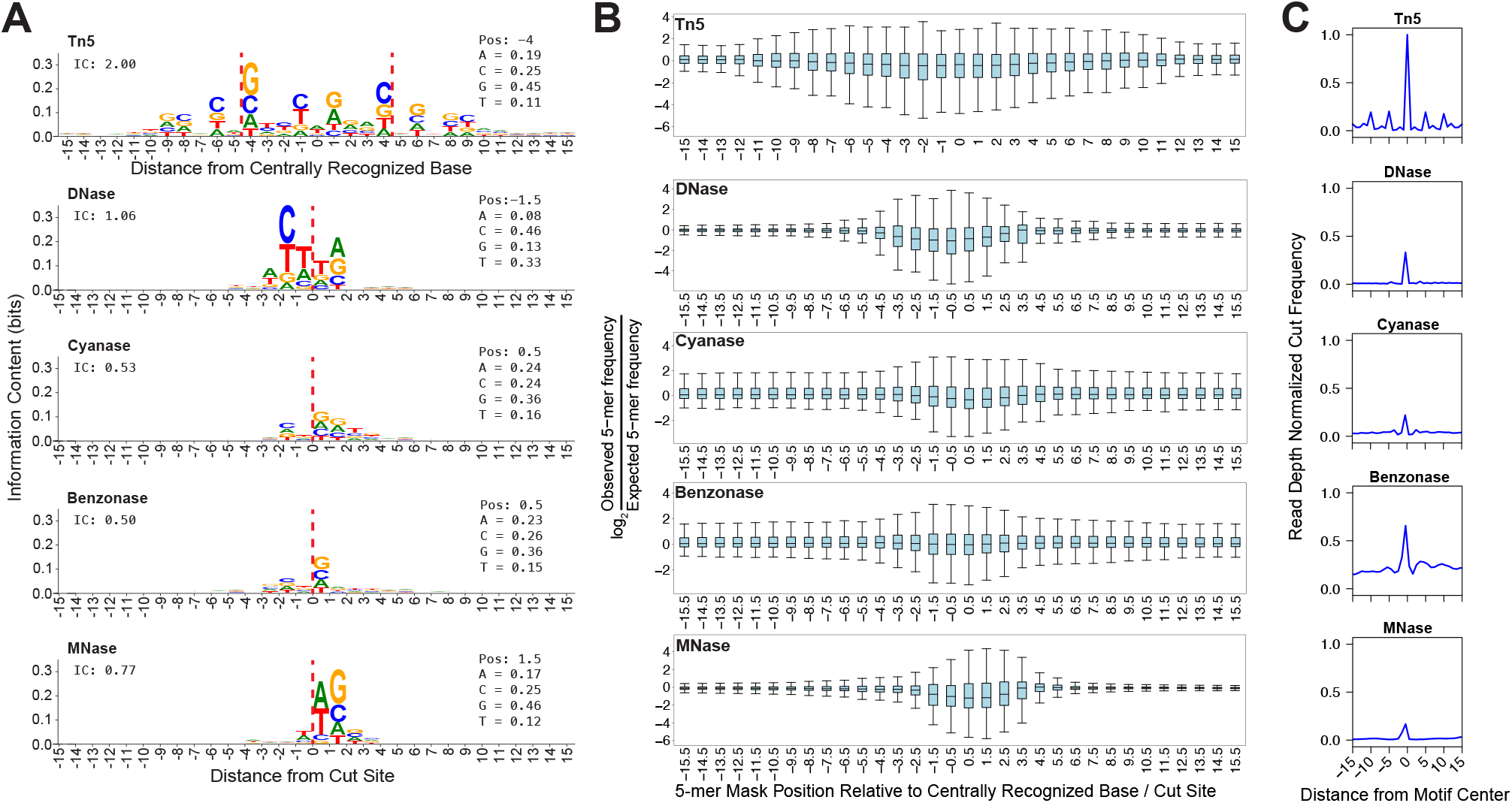
Tn5 sequence bias is more complex than nuclease sequence bias. A) The seqLogo sequence bias motifs are corrected for background nucleotide content and they illustrate that Tn5 bias is wider and more complex than other commonly used chromatin accessibility enzymes. Nucleotide frequencies listed in the inset to the right correspond to the highest information content position; for instance, G is found at position -4 45% of the instances that Tn5 inserts into DNA. Nuclease cleavage sites are indicated by dashed red line. Total information content (IC) from positions -10 to 10 is listed in the inset to the upper left. B) We plotted the log_2_ (observed/expected 5-mer frequency) for all positions surrounding DNA cleavage sites or Tn5 recognition sites as box and whisker plots. C) We plotted read depth normalized composite signal from various molecular genomics assays for the top 400,000 sites that conform most stringently to the respective enzyme’s bias motif.

If nucleotide sequence far from the Tn5 recognition site center influences cleavage and insertion, then we would expect that k-mers found at positions distal from Tn5’s centrally recognized base would deviate substantially from random expectation. We quantified the influence of distal and proximal k-mers on cleavage bias by plotting the log_2_ ratio of observed 5-mer frequency to expected 5-mer frequency for all 1024 5-mers; expected frequency is based on the genomic frequency of each 5-mer. The log_2_(observed/expected) ratio is zero if a 5-mer is observed at the expected frequency. We find that the distribution of this ratio becomes more tightly centered around zero as we query more distal 5-mers (Figure 1B). The distribution does not stabilize for Tn5 until we exceed 11 bases from the Tn5 recognition site, while the other four enzymes stabilize within 4 bases of the cleavage event. This analysis confirms that distal sequences more strongly influence Tn5-mediated cleavage compared to the other enzymes.

The background corrected sequence motif (Figure 1A) for Tn5 is a reverse palindrome, which is common for homodimeric DNA binding factors (Sasse et al. 2017; Welboren et al. 2009). The consensus Tn5 recognition sequence contains CAG trimers present on each strand of the DNA, each CAG is displaced from the adjacent reverse complemented CAG by 5 bases. This unique feature suggested that Tn5 dimers may interact with this sequence motif in multiple orientations to direct the transposition of adapters with higher frequencies. We plotted the read-depth normalized composite profile of each enzyme’s bias motif to compare their relative strength of directing DNA cleavage (Figure 1C). The Tn5 recognition site directs a central peak that is substantially higher than the other enzymes. The composite profiles also highlights peaks 5 and 10 base pairs downstream and upstream of the centrally recognized base (Figure 1C). These flanking peaks are not observed in the other nucleases, although they are comparable in intensity as the central peak of other enzymes. Although not definitive, this pattern is consistent with Tn5 dimers interacting with opposite sides of the DNA helix, displaced by five bases. Taken together, the length of the Tn5 bias motif, the strong influence of distal k-mers, the stronger cutting preference, and 5-base spaced cutting pattern all preclude conventional k-mer based approaches for correcting Tn5 sequence bias.

### Tn5 sequence bias is complex and not modeled well by previous methods

A key assumption of representing sequence biases as probability matrices or seqLogos is that any nucleotide observed at a position within the motif does not influence the probability of observing nucleotides in other positions, this property is referred to as positional independence (Guertin et al. 2012; Sharon et al. 2008). If each position is independent from the others, observed k-mer frequencies should be equal to those expected based on nucleotide frequency in the position probability matrix. To test Tn5 sequence bias position independence, we determined the observed frequency of each 3-mer found in position -6 to -4 relative to the central Tn5 recognition base and divided by the expected frequency (Figure 2A). We find that 3-mers are observed at frequencies that deviate from the calculated expectation. We can infer basic features describing which 3-mers are disfavored (NAA) and preferred (NCA), but characterizing a comprehensive set of basic rules for many k-mer combinations and k-mer positions is not straightforward. However, since k-mer scaling does consider dependencies between positions, k-mer scaling remains an attractive starting point for developing more complex bias correction models.

**Fig. 2.**
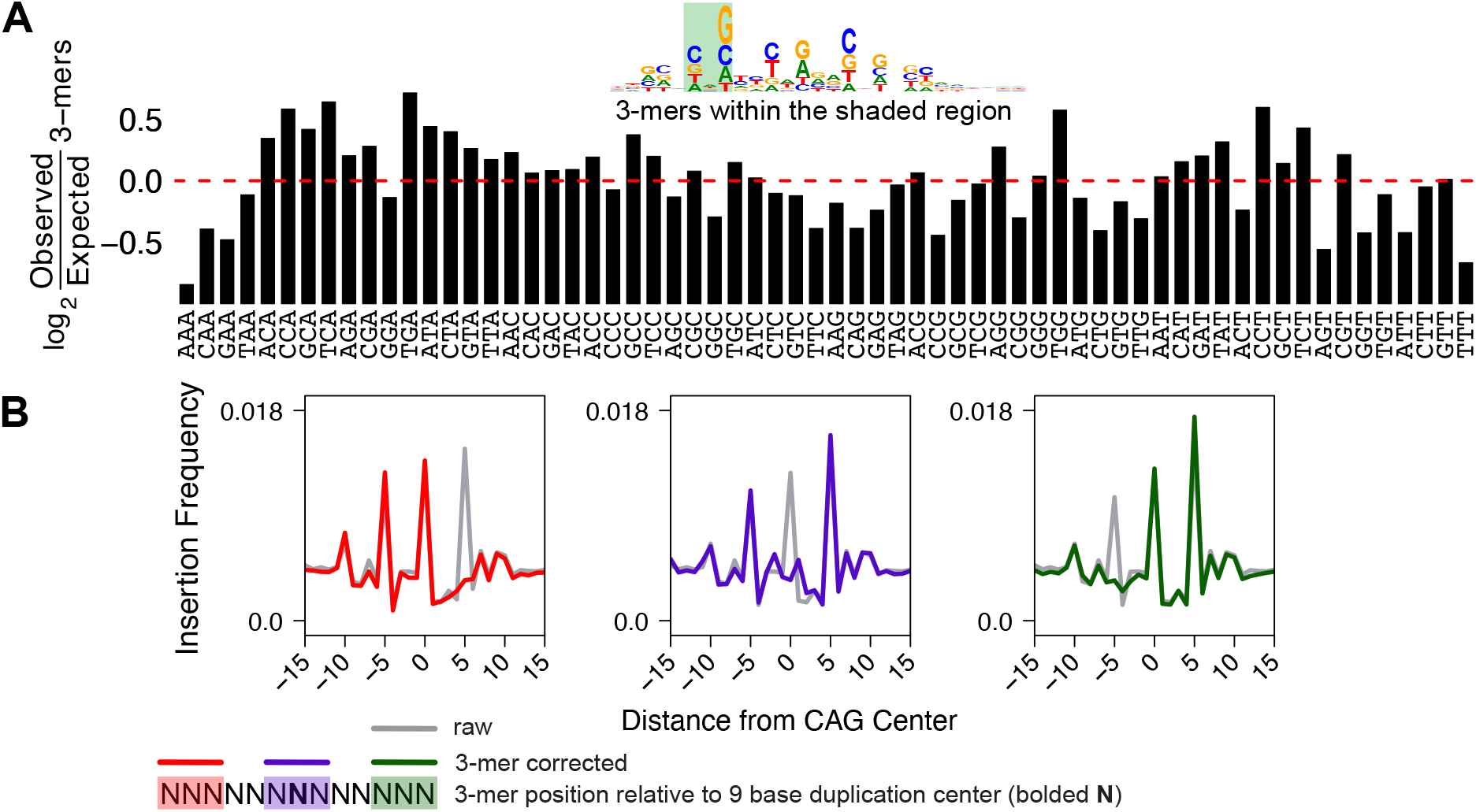
K-mers capture more bias complexity than weight matrices and provide a basis for which to build more sophisticated bias-correction models. A) We quantified the frequency that all 64 3-mers occur at positions -4 to -6 from the center of a Tn5 recognition site and compared observed frequency to the position weight matrix prediction. A bar chart of log_2_ observed divided by expected (from k-mer prevalence in the bias position probability matrix) k-mer frequency highlights the interdependency between sequences are different positions. B) We plotted the signal from 400,000 randomly selected “CAG” instances in the genome. Overlaid on top of the unscaled signal, each plot shows that the individual peaks from composite profile can be corrected by rationally designed spaced k-mer scaling, based on the position of the scaling k-mer and the peak.

Next we tested the feasibility of using rationally spaced k-mers to systematically reduce Tn5 bias. The 3-mer “CAG” is over represented in three positions in the Tn5 bias and each CAG bias is displaced by 5 bases (Figure 1A). We measured the composite ATAC-seq signal in the window centered on 400,000 random CAG instances in the hg38 genome assembly. As expected, we observe that the CAG 3-mer directs cutting in three peaks, each displaced from one another by 5 bases (Figure 2B). Conventional k-mer scaling is performed with any reasonably short k-mer size (<9 bases) and the k-mer can be positioned at any distance from the observed Tn5 recognition site. In an effort to determine whether k-mer scaling would effectively flatten these three peaks, we used `seqOutBias` to scale reads using 3-mers that are located at the Tn5 recognition site and up/down stream by 5 bases. Recall that each peak corresponds to a peak of Tn5 recognition and that each peak is relative to the CAG 3-mer that anchors the plot at position x = 0. Therefore, we expect that scaling based on the k-mer located 5 bases upstream will flatten the +5 peak because the CAG directing this downstream peak is located 5 bases upstream. Likewise, the central k-mer scaling would abolish the central peak and downstream k-mer scaling would ablate the upstream peak. This exercise indicates that we can rationally design k-mer scaling to correct known biases, but it also highlights the fact that multiple kmers are needed to correct biases even within the context of this simple example. Therefore, we pursued a rule ensemble modeling approach to integrate multiple k-mer sizes from a range of positions relative to the Tn5 recognition site as input data and determine whether interactions between these inputs contribute to Tn5 insertion bias.

### Rule ensemble modeling leverages interaction terms and k-mer scaling to correct sequence biases

We previously developed `seqOutBias`, which directly scales chromatin accessibility data by expected/observed k-mer frequency. Although we specify `seqOutBias` to scale based on a 6-mer centered on the DNA cleavage event for DNase, one can specify any k-mer length and relative k-mer position. Our data indicate that Tn5 bias is too broad to scale with a single k-mer and this suggests that bias correction would need to incorporate information from several k-mer positions. Therefore, we chose to use `seqOutBias` scaling outputs with different invocations of k-mer positions and lengths as the input for the rule ensemble model, and to test its capacity to correct Tn5 bias (Figure 3A). Sequence biases were first characterized by quantifying enzymatic cut frequency in average composite profiles that are centered on transcription factor binding motifs (He et al. 2014; Sung et al. 2014). We previously developed a model (seqToSign) that reproduced these composite profiles by scaling the genome-wide cut frequency of each k-mer found in a given position in the composite by the number of occurrences of the k-mer at that position (Sung et al. 2014). We attempted a similar approach to predict biased composites by using the inverse of `seqOutBias` scale factors as an input, since this value represents the genomic cut frequency for a k-mer. For each k-mer we multiplied its frequency at each position in the composite by the inverse of that k-mer’s corresponding `seqOutBias` scaling factor. We summed these values for all 4*k* k-mers at each position in the composite profile (Figure 3B). Each of these values at each position in the composite is a covariate input for the rule ensemble model and we repeated the process for many k-mer length and relative k-mer position combinations (Figure S3A). We trained a rule ensemble model using these covariate inputs to predict multiple transcription factor composite profiles. The output of this trained model is the formulaic combination of k-mer positions using linear regression coefficients and decision trees (Figure 3C). To test the efficacy of the model, we first combined the scaled read values from each k-mer covariate according to the output formula. This generated a single rule ensemble-scaled read file which we tested for its ability to correct test group composite profiles to resemble theoretically unbiased genomic cleavage (Figure 3D).

**Fig. 3.**
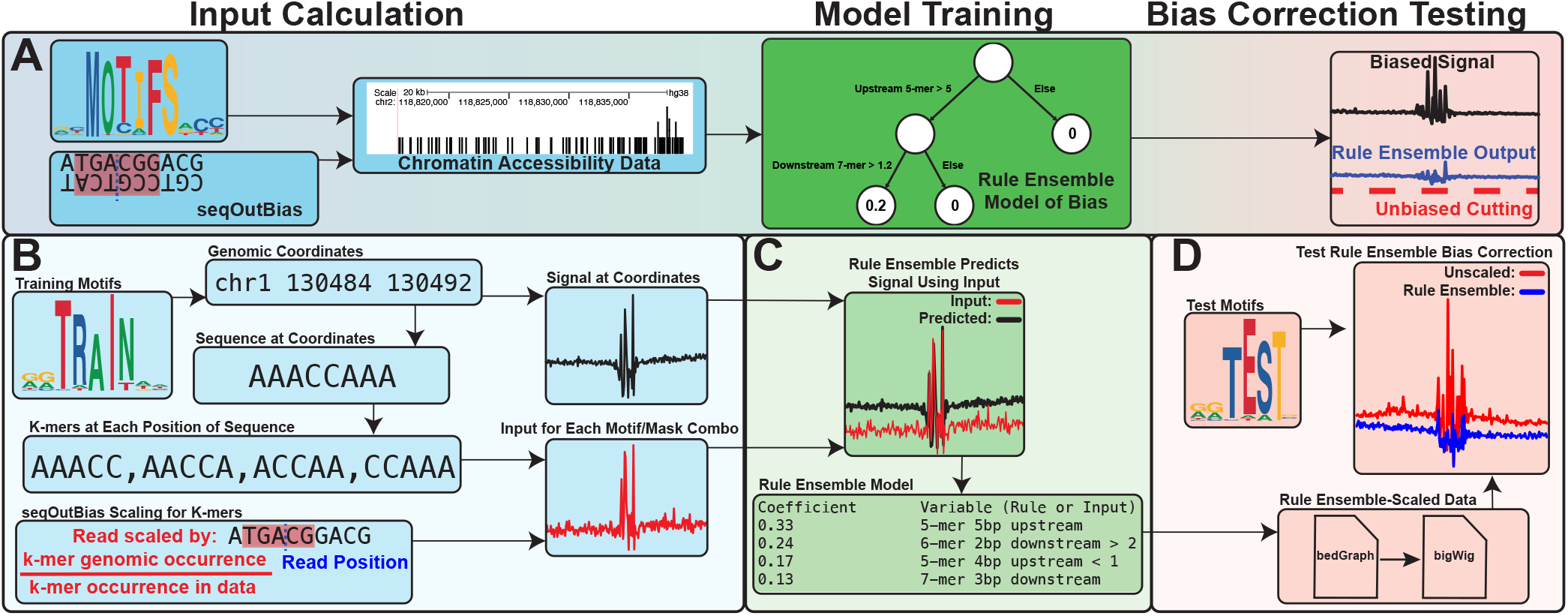
Rule ensemble modeling of enzymatic bias combines k-mer scaling approaches to enhance bias correction. A) We generate the rule ensemble input by combining `seqOutBias` scaling output with k-mer frequencies observed at transcription factor motifs found throughout the genome. This input is then used to train a rule ensemble model to predict the enzymatic sequence bias. Finally, the model is tested for its ability to correct this bias. B) For each training motif, we identify the 400,000 occurrences in the genome that conform most stringently to the weight matrix. We extract sequences (200 bases) in the region for each identified motif and calculate the frequency of each k-mer at each position relative to the motif center. We perform independent runs of `seqOutBias` to calculate scale factors for each k-mer in each position within a defined window from the center of the motif. K-mer frequency for each position is then multiplied by the inverse scale factor for every frame of reference. These are the rule ensemble input values for each motif/k-mer (size and position) pairing. C) We train a rule ensemble model to predict the biased signal measured at each of these region sets in a deproteinized data set. D) Output from `seqOutBias` corresponding to all input frames of reference are then combined according to the rule ensemble model to generate a single BED or bigWig file with bias-corrected values for each sequence read. Successful bias correction is evaluated using a held-out test set of TF motifs.

### Choosing training and test transcription factor motifs

Transcription factors recognize a diversity of DNA sequences that vary by length, degeneracy, and nucleotide content. We chose training and test data for the model by selecting sequence motifs based on these features (Figure 4A). In a sequence logo representation of DNA binding preference, the information content is a function of sequence length and degeneracy. We used hierarchical clustering of these values for 43 input motifs that represent a wide range of information content and GC content, encompassing the majority of known motifs (Figure 4B). Groupings consisted of either doublets or triplets, with a single singlet. Within a group, a single motif was assigned as testing data and the others were assigned as training data.

**Fig. 4.**
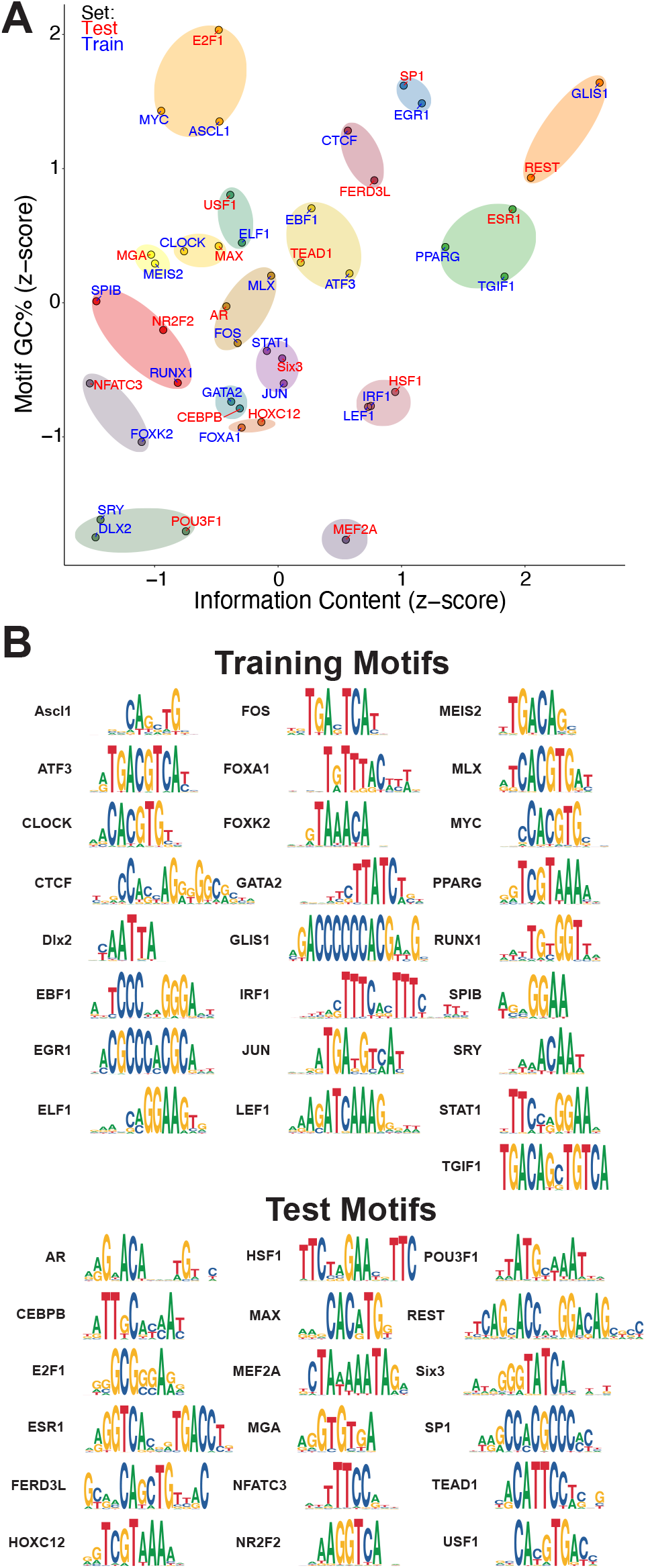
Clustering of transcription factor motifs creates diverse test and training sets. A) We clustered transcription factor motifs into test and training sets based on motif information content and GC content. We indicate clusters with transparent ovals. Training and test sets are indicated as red or blue text that indicates the motif name. B) The seqLogos of training and test transcription factor motifs illustrate their diversity.

### Rule ensemble modeling corrects DNase sequence bias

We chose rule ensemble modeling because the importance of covariate inputs are easily interpreted in the overall model and for any individual model prediction. Since DNase-seq biases are well-characterized, we have an expectation of the importance of positions that lead to DNase specificity. We chose to test the feasibility of this model using naked DNA DNase-seq data. We invoked `seqOutBias` with staggered 5-mers within 9.5 base pairs of the cut site as input for the model. The DNase rule ensemble model confirmed that the 3 bases on either side of the cleavage event most strongly influence DNase cleavage (Figure 5A).

**Fig. 5.**
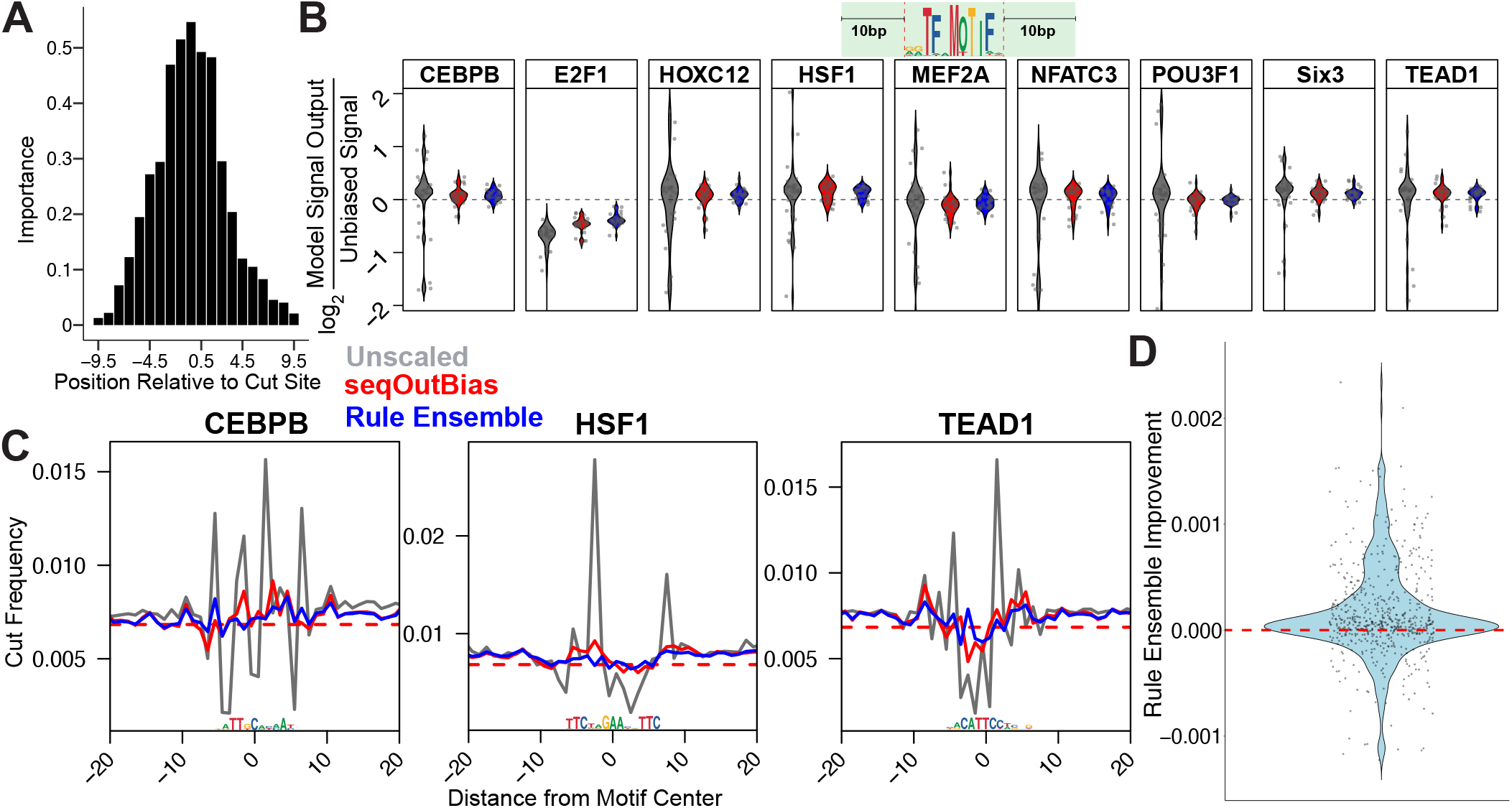
Rule ensemble modeling effectively corrects DNase bias. A) Rule ensemble modeling provides information about the importance of variable in the model. The positions proximal to DNase cleavage are most important. B) Violin plots quantify the log_2_ ratio of unbiased signal to output for each transcription factor’s composite profile given the method of k-mer scaling. Regions measured in each plot are within ±10 base pairs of each motif, as indicated by the graphic above the figure panel. C) We plotted composite profiles of transcription factors from rule ensemble output compared with direct k-mer scaling of the 5-mer that encompasses the five most influential positions in panel A. D) We visualized improvement of rule ensemble modeling over seqOutBias by using a violin plot of distance from calculated random cut frequency for rule ensemble output subtracted from seqOutBias. Positions which rule ensemble outperforms `seqOutBias` will be above 0 (dashed red line); 68% of positions in this plot were improved.

We visualized the correction of single nucleotide bias by calculating the log_2_ ratio of theoretically unbiased signal to either unscaled, direct k-mer scaling (seqOutBias), or rule ensemble output within the region of ±10 base pairs for each motif (Figure 5B & Figure S4A). Our results indicate that direct k-mer scaling and rule ensemble modeling correct DNase bias comparably well. This is most easily illustrated by observing individual composite profiles (Figure 5C & Figure S4B). Although improvement is comparable between direct k-mer scaling and rule ensemble modeling, we sought to directly compare the two more rigorously. We calculated the absolute value of the difference between each corrected value and random cleavage for each point within 10 base pairs of each motif for both bias correction methods. This metric represents how much the values deviate from perfect bias correction for each method (`seqOutBias` and rule ensemble). For each position, we subtracted the rule ensemble deviation from unbiased cleavage value from the seqOutBias deviation from unbiased cleavage and plotted the difference (Figure 5D). Each point represents the difference in improvement between `seqOutBias` and rule ensemble modeling. For each point greater than 0, the rule ensemble modeling outperforms `seqOutBias` (Figure 5D). We find that 68% of all points showed improvement, indicating that rule ensemble modeling generally outperforms direct k-mer scaling for correcting DNase bias.

### Rule ensemble modeling corrects local Tn5 transposition bias

Since this modeling approach corrected DNase bias, we developed a rule ensemble model to correct bias from deproteinated ATAC-seq data. Since Tn5 bias is more complex than DNase, we ran `seqOutBias` for all contiguous 5-mers, 6-mers, 7-mers, and spaced 6-mers within 19 bases of the DNA cleavage/insertion site for the model input. In total, we included 663 distinct k-mer combinations as inputs (Figure S3A). The number of input variables was much higher compared to DNase, so we prioritized the input covariates by first modeling the data using linear regression to determine the most influential k-mers, then we incorporated the most influential 10% of these variables into a full rule ensemble model to predict the biased signal for each training motif (Figure 3C). The relative importance values for each nucleotide position revealed three central peaks that are separated by 5 bases (Figure 6A). Comparison of individual composite profile traces highlights the improvements (Figure 6B & Figure S5A&B). We systematically measured how bias-corrected signal deviated from random DNA cleavage at sequence motifs (±10 bases) and compared to unscaled and k-mer scaled correction (Figure 6C). The rule ensemble bias correction globally outperforms k-mer scaling. For each individual position, the rule ensemble model outperforms k-mer scaling 78% of the time (Figure 6D). Examination of the 22% of positions with worse performance reveals the magnitude by which the model is outperformed is modest compared to the magnitude of improvement for the other 78% of instances. This can be visualized by comparing the distribution above zero (improvement) versus below zero (outperformed by k-mer) in Figure 6D. Visual inspection of the overlay traces of the 18 test motif composite profiles illustrates that rule ensemble models effectively flatten these profiles (Figure 6E & FigureS5B). The traces approach the theoretical random cleavage red dotted line more closely than k-mer scaling alone (Figure 6E & S5B). Therefore, the local sequence bias, which we define as the region within 10 bases of sequence motifs, is more effectively corrected with rule ensemble modeling.

**Fig. 6.**
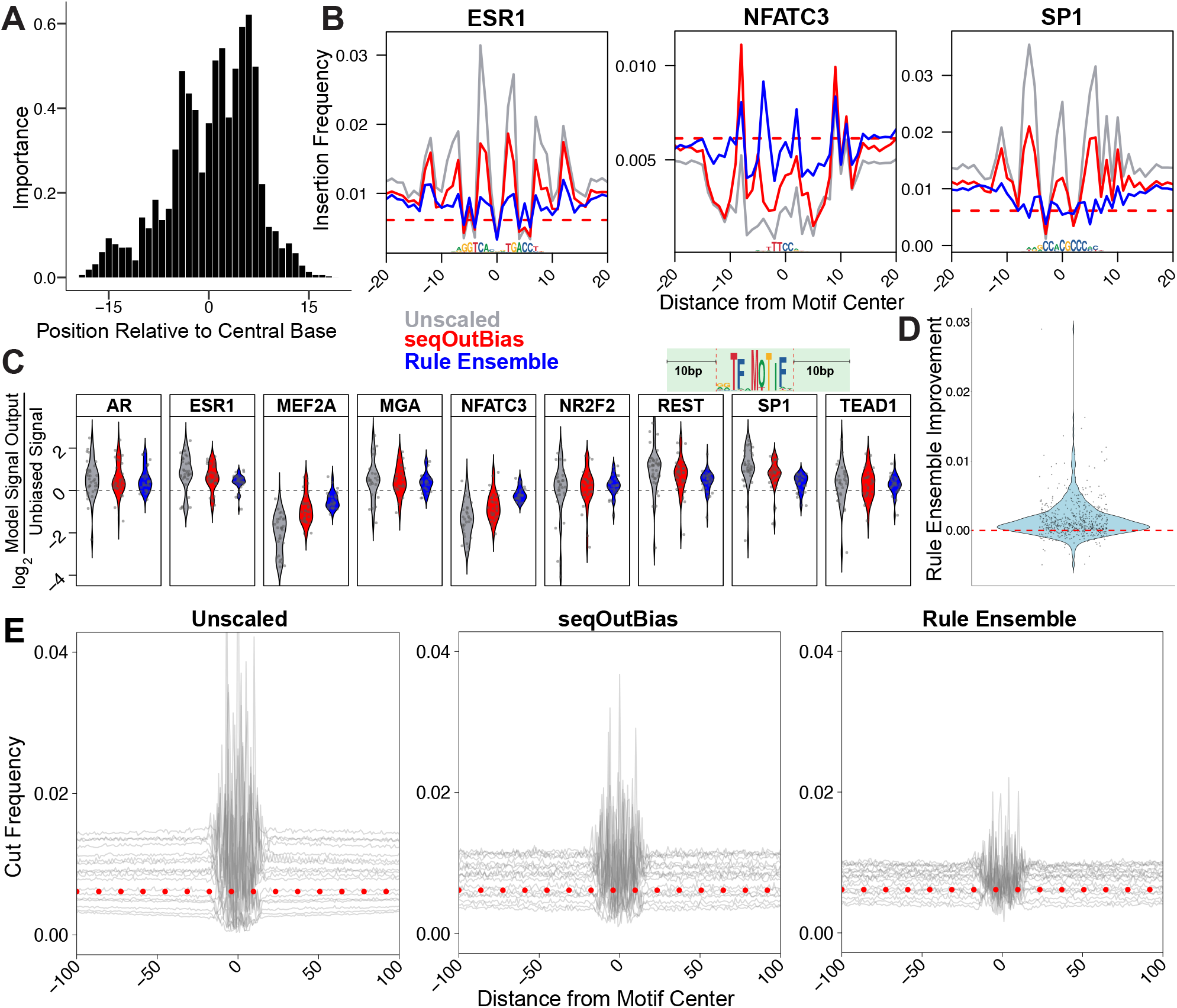
Rule ensemble modeling corrects Tn5 transposon single nucleotide bias in ATAC-seq data. A) The importance values for each position relative to Tn5’s recognition site mirror the information content values of positions in the seqLogo representation of Tn5 bias. B) We directly compare bias correction of Tn5 data by `seqOutBias` and rule ensemble modeling by visualizing the composite corrected signals and comparing to the unscaled composite traces. C) We quantify the single nucleotide correction for each transcription factor’s composite profile by measuring divergence from unbiased signal. Regions measured are within ±10 base pairs of each motif, as indicated by the graphic above the figure panel. D) We visualized improvement of rule ensemble modeling over `seqOutBias` by using a violin plot of distance from calculated random cut frequency for rule ensemble output subtracted from `seqOutBias`. Positions which rule ensemble outperforms `seqOutBias` will be above 0 (dashed red line); 78% of positions in this plot were improved and those that are not improved have lower magnitude deviations from the expectation of random cleavage. E) Composite profile overlays of test motifs highlight the improvement of rule ensemble modeling compared to `seqOutBias` correction. The red dashed line indicates the calculated random cleavage frequency.

### Rule ensemble modeling corrects regional sequence biases caused by GC-content

We developed these models and measured correction of enzyme biases using these composite motif profiles because this visual representation is an intuitive way to observe enzyme biases in genomic assays. As we move away from the sequence “anchor” in DNase-seq composite profiles, the traces flatten and approach the expectation of random cleavage because the sequence content in these regions is more random (Figure 7A). Unlike DNase-seq, the composite profiles for ATAC-seq do not approach random expectation, even at distances 100 bases from the sequence motif center. The Tn5 recognition site is GC-rich (Figure 1A) and it is known that AT/GC richness is clustered throughout the genome (International Human Genome Sequencing Consortium 2001). We hypothesized that Tn5 preference for GC-rich regions would lead to generally elevated signal within these regions, accompanied by depleted signal in AT-rich regions. Therefore, we determined whether there is a relationship between motif GC content and this regional “baseline signal”. Motif GC content linearly correlates with baseline signal (Figure 7B&C). The rule ensemble modeling outperforms `seqOutBias` for 88% of the positions when regional GC bias correction (Figure 7D) and scaling within the motif (Figure 6D) are considered together. Rule ensemble modeling effectively corrects Tn5’s regional and local sequence biases.

**Fig. 7.**
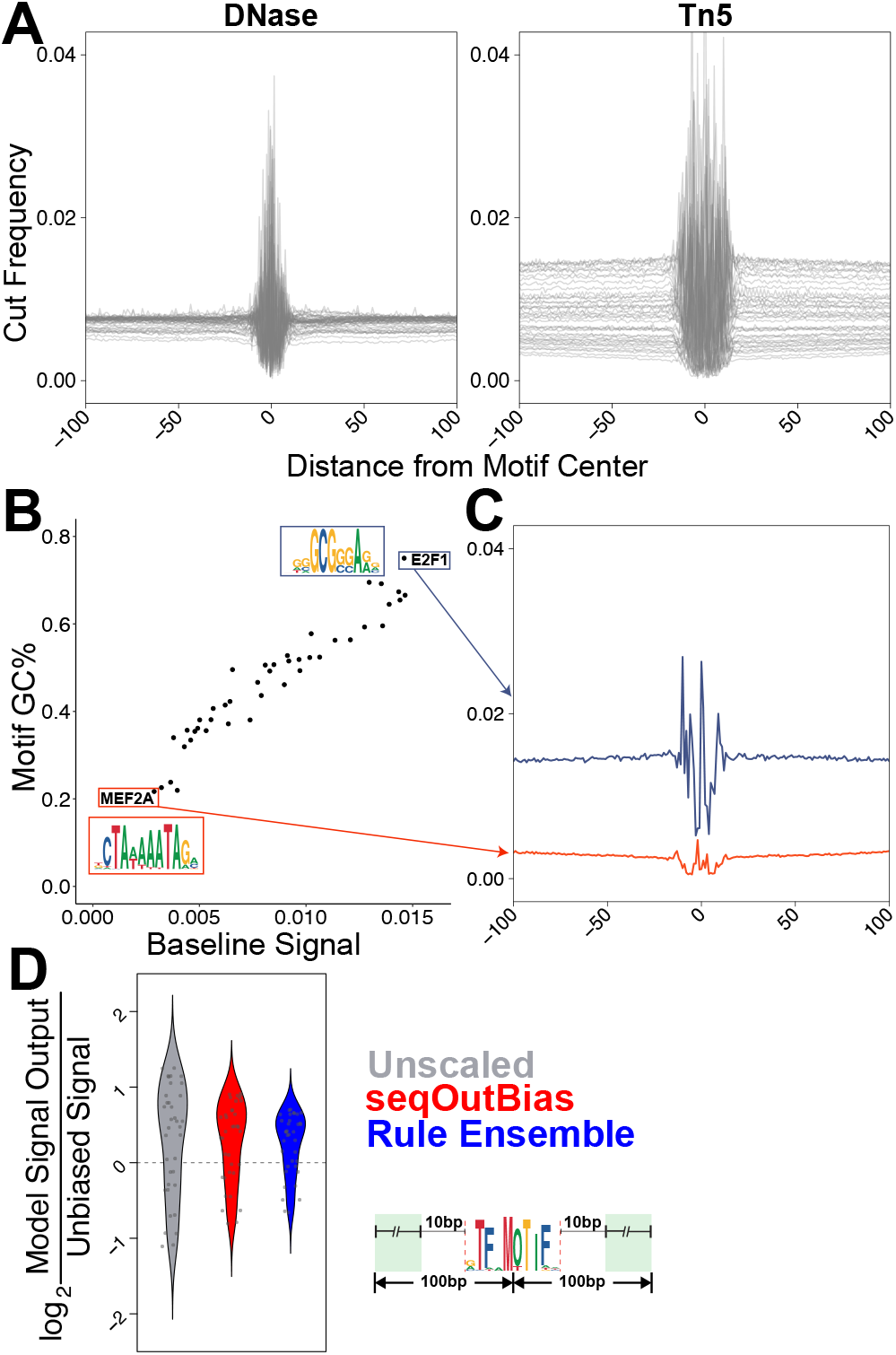
Rule ensemble modeling corrects regional sequence biases caused by GC-content. A) We illustrate the range of baselines for each enzyme by plotting unscaled overlays of DNase and Tn5 composite profiles for the 43 transcription factors surveyed. B) A plot of motif GC content vs. baseline signal indicates a linear and correlated relationship between the variables. C) The E2F1 and MEF2A motifs have very different GC content and this is reflected in their baseline signal within the composite traces. D) Baseline correction is measured by the ratio of unscaled, seqOutBias, and rule ensemble scaled reads to random cleavage. Two points for each motif are plotted; each point is the average of upstream or downstream signal in the window ±10 base pairs from the edge of the motif spanning 100 bases from the motif center (green box).

## Discussion

Transcription factor binding sites and promoters are the most interesting regions of the genome with respect to gene regulation. ATAC-seq signal correlates with the regulatory potential of these regions. ATAC-seq is a simple experimental assay, but analysis of the data requires dedicated pipelines, specialized software, and unique considerations (Smith et al. 2021). Importantly, Tn5 sequence bias was described in the first ATAC-seq paper (Buenrostro et al. 2013) and many groups have developed methods to characterize and correct these biases. Approaches often combine bias correction and footprinting or propose models that cannot be easily interpreted. Here, we provide a workflow that directly scales individual ATAC-seq reads to correct Tn5 bias and provides common output files that can be used for peak calling or footprinting. The rule ensemble modeling approach that we employed is interpretable globally, in that we identify the positions relative to Tn5 recognition that influence the model. Moreover, for any individual scaled read the model is interpretable locally. If a rule does not apply to the local sequence read, then the rule drops out of the model and does not contribute to that individual instance of read scaling.

Enzymatic sequence preferences must be considered when interpreting molecular genomics data, particularly when the data is single-nucleotide resolution. We introduce this rule ensemble framework as a novel method that efficiently scales individual sequence reads to correct Tn5 bias.

## Methods

### Reproducible analysis and plotting code

We provide a detailed and reproducible vignette to reproduce all analyses and figure panels directly from raw data here: https://github.com/guertinlab/Tn5bias/tree/master/Manuscript_Vignette. We provide a workflow vignette using only chr21 ATAC-seq reads, the chr21 reference genome, and estrogen receptor motifs as a quick-start reference document: https://github.com/guertinlab/Tn5bias/tree/master/seqOutATACBias_workflow_Vignette

### Chromatin accessibility data preprocessing and read alignment

We downloaded the hg38 and mm39 reference genomes from the UCSC genome browser (Consortium 2002; International Human Genome Sequencing Consortium 2001). We retrieved data sets for the respective nucleases and Tn5 from the NCBI SRA (sequence read archive), in fastq format, using `fasterq-dump` (Leinonen et al. 2010). The following SRA accession numbers were used for mouse (mm39) liver data: SRR535737, SRR535738, SRR535739, SRR535740, SRR535741, SRR535742, SRR535743, SRR535744 (Grøntved et al. 2012), SRR5723785 (Iwata-Otsubo et al. 2017). The following SRA accession numbers were used for human (hg38) DNase (lung fibroblast) SRR769954 (Lazarovici et al. 2013) and Tn5 (T lymphoblast) SRR5123141 (Martins et al. 2018). Reads from each data set were aligned to their respective reference genomes using `bowtie2` (Langmead and Salzberg 2012), then sorted and converted to BAM files with `samtools` (Danecek et al. 2021).

### Determining sequences around each cleavage site

All aligned reads (plus, minus, and unseparated) were used individually as input for unscaled `seqOutBias` (Martins et al. 2018) runs, which generated bigWig and bed format output files. We deconvolved independent reads that aligned to the same genomic coordinate into separate bed file entries, then converted to sequence files using `fastaFromBed` (Quinlan and Hall 2010). Sequences corresponding to minus-aligned reads were reverse complemented before concatenating them with the sequences corresponding to plus aligned reads.

### Counting positional nucleotide frequencies for each enzyme

We determined nucleotide counts at each position relative to the start of a read by loading all the sequences into R, separating sequences into columns, and tallying nucleotide bases at each position. Results were output in TRANSFAC format and input into `Weblogo` (Gavin E. Crooks and Brenner 2004).

### Plotting background nucleotide frequency-corrected sequence logos

We desired background corrected information content values for our sequence logos generated from `Weblogo` (Gavin E. Crooks and Brenner 2004). We retrieved these values by modifying the source code of the `Weblogo` command line interface Python. A step-by-step guide on the modifications we made to retrieve these values is included in the vignette on Github. As a coherence check, we calculated the background corrected information content values for each position. First, we calculated the Shannon entropy:

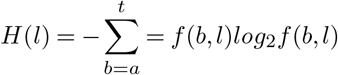

Here, H(l) is the entropy at any given position, and f(b,l) is the frequency of a base (b) at this position (l). Subtracting this value from 2 is the classic calculation for information content. We next calculated the background entropy:

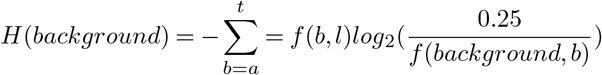

Where H(background) is the correction for background nucleotide frequency. And f(background,b) is the background frequency (background) for a base (b) Thus, for any position the background corrected information content is:

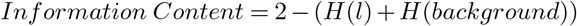

These values were plotted using the `Weblogo` command line interface.

### Determining enzyme bias motif genomic coordinates

Starting with the TRANSFAC format nucleotide count files, we used `transfac2meme` (Bailey et al. 2015) to convert to the meme file format. We then used these files as input into `FIMO` to generate region sets for each bias motif using the appropriate refererence genome (Grant et al. 2011). The highest scoring 400,000 regions were used for each composite profile plot.

### Plotting composite signal of genomic coordinates

We visualized the average signal at genomic coordinates by aligning plus and minus locations on the same position and retrieving the signal in the plotted interval from a given data set’s bigWig file, using the `bigWig` R package (Martins 2014). We then divided all signal in this interval by the number of genomic coordinates plotted, to calculate the average cut or insertion (for Tn5) frequency at this position relative to the motif.

### Determining background nucleotide frequencies of reference genomes

Background nucleotide frequencies for reference genomes were calculated by using the grep command to count the occurrences for a given nucleotide.

### Calculating scale factors for k-mer position

Genomic and data set k-mer frequencies were determined by using the `seqOutBias table` (Martins et al. 2018) command to tally each k-mer in the reference genome and those in the data sets for each k-mer size and position. We then used these k-mer counts to calculate scale factors for each k-mer by dividing the expected k-mer frequency in the genome by the observed k-mer frequency in the data set (Martins et al. 2018).

### Calculating observed and expected upstream k-mers

Starting with the sequences flanking each insertion in the Tn5 data set in fasta format, we counted each 3-mer in relevant positions from the cut site to determine observed k-mer occurrence. To determine expected k-mer occurrence, nucleotide frequency from the sequence logo motif at each position was multiplied together, in all possible combinations, to construct each possible k-mer. This value is the expected frequency for each k-mer.; we plotted the log_2_ ratio of observed to expected k-mers.

### CAG peak direction

We generated a list of all 3-mer locations in the genome by invoking the `seqOutBias dump` (Martins et al. 2018) command using a 3-mer .tbl file (output from previous invocations of `seqOutBias table`) for input. From this file, we determined the genomic location of all CAG instances in the reference genome. We then plotted ATAC signal at 400,000 random CAG instances from this bed file. Rationally designed masks were implemented using the `seqOutBias` (Martins et al. 2018) software with the noted k-mer masks.

### Transcription factor motif genomic interval determinations

Motifs for each transcription factor included in the test and training sets were downloaded from the `JASPAR` (Castro-Mondragon et al. 2022) database in meme format. These motifs were then used as input, along with the hg38 reference genome, into `FIMO` to output genomic regions conforming best to the motif. We took the top 400,000 highest scoring genomic instances for each motif for both the plus and minus strands and used them as input for the rule ensemble model.

### Rule ensemble target input

We calculated the target values by plotting the composite signal at each transcription factor’s genomic coordinates using unscaled data produced from `seqOutBias` (Martins et al. 2018), using the `-no-scale` option. `bigWig` files were accessed and plotted using the `bigWig` R package (Martins 2014). These plotted values were normalized using a common factor, to preserve variation between motifs and allow for accurate rule ensemble prediction.

### K-mer frequency calculation

We determined k-mer frequency using the 400,000 genomic locations for each strand of each transcription factor motif previously generated using `FIMO`. From these locations, we retrieved the sequences using `fastaFromBed` (Quinlan and Hall 2010). We split each sequence into k-sized slices for each k-mer. For example, using a 5-mer, the sequence “AAACCAAA” would be split into: AAACC,AACCA,ACCAA,CCAAA. Each of these sections are a position within the original sequence: position 1-AAACC; position 2-AACCA; etc. For each position, we then determined the frequency of each k-mer among the original 400,000 input sequences.

### Rule ensemble independent variable input

We computed the rule ensemble input for each combination of transcription factor motif region set and k-mer size/position. As we previously determined k-mer frequency at each position of a composite profile, we multiply these frequencies by the inverse of the scale factor for each matching k-mer, which was also described above. Finally, we sum these values for all possible k-mers at a position for the input value. This process was repeated for each modeled position in the composite corresponding to each included k-mer size and location relative to the cut site. These values are the `seqOutBias` predicted values output for a k-mer size and position, and modeled genomic regions.

### Rule ensemble modeling

After calculating the independent variable input, a rule ensemble model was trained to predict the training set biased target values using the `prediction rule ensemble` package in R (Fokkema 2017). This package implements the original rule ensemble modeling framework (Friedman and Popescu 2008). The output rules and scaling coefficients were recorded as a single equation for later implementation.

### Rule ensemble model implementation

We implemented the rule ensemble model by combining scaled `seqOutBias` (Martins et al. 2018) output as defined by the model. We first aggregated the required `seqOutBias` output by combining output bed format files using `unionbedg` software (Quinlan and Hall 2010). These output values were then combined by applying the modeling equation at every read to generate a rule ensemble-scaled bed file. This bed file was then scaled to the same total read depth as the original, unscaled data. Finally, we converted the bed file to bigWig format for subsequent analysis using the `bedGraphToBigWig` command line interface (Kent et al. 2010).

### Calculating positional importances from rule ensemble models

We calculated variable importances, for each position within our range of inputs to visualize how the rule ensemble model combined input positions. An input contributed to the importance of a position if it used that position to model k-mer bias. We first determined total importance for each position by adding together the importances of each input which included a given position. This was how we calculated importances for our rule ensemble DNase model. The Tn5 rule ensemble model did not have equal positional coverage for all inputs, so we normalized all total position importance by the number of inputs that included the respective position in the input masks.

### Calculating improvement of rule ensemble modeling compared to seqOutBias

We directly compared the improvement of rule ensemble modeling over `seqOutBias` by first determining the absolute value difference between scaled output from either method and the calculated unbiased output. This value was determined for each position in our test set composite profiles. Each of these values is the disparity between either method and perfect bias correction at a given position—a value of 0 means the method perfectly corrected bias at this position. We then took these values and for each position, subtracted the rule ensemble output from the `seqOutBias` output. For every value above 0, rule ensemble modeling outperforms `seqOutBias` correction.

## ACKNOWLEDGEMENTS

This work was funded by R35-GM128635 to MJG. A Science Post Graduate Scholarship Fund award from the Bureau of Indian Education supported JBW. We thank Sathyan Mattada, Thomas Scott, Arun Dutta, and Abigail Wolpe for critical feedback.

## AUTHOR CONTRIBUTIONS

JBW and MJG analyzed the data. ALM and MJG updated `seqOutBias` to accommodate ATAC data. JBW and MJG conceptualized and developed the project. JBW and MJG wrote the manuscript.

---

**Fig. S1.**
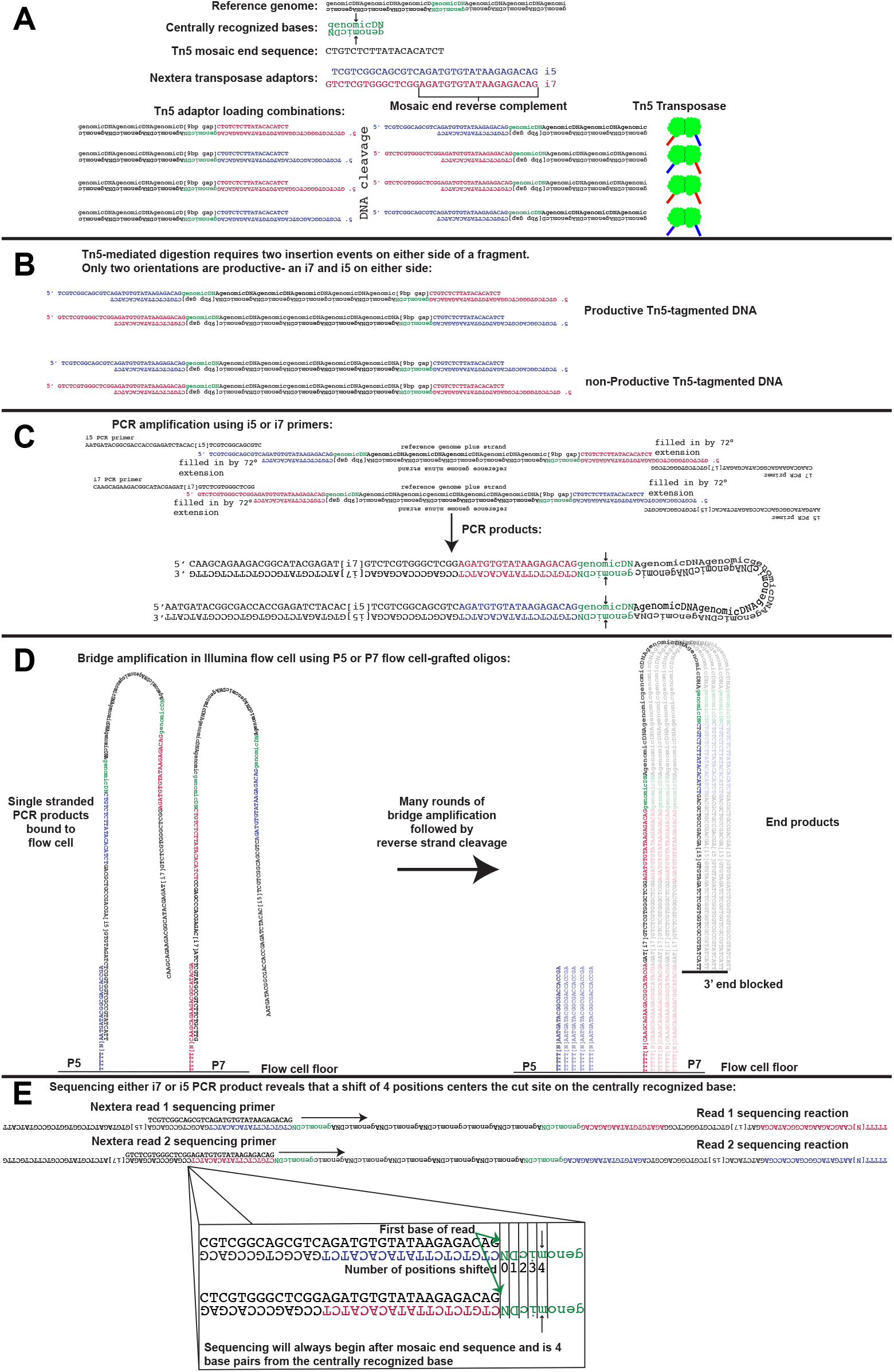
Shifting forward and reverse aligned ATAC-seq data reads by 4 bases captures the center of the Tn5 duplication. A) Each monomer of the Tn5 transposase dimer is loaded with one of two Nextera adapter sequences. During an insertion event, each adapter is attached to one side of the duplicated 9 base pairs. Each adapter sequence is a concatenation of the Tn5 hyperactive mosaic end sequence and either an i7 or i5 sequencing primer. Each Tn5 dimer is capable of four possible DNA insertion events, based on which adapters are loaded into each Tn5 monomer. B) For a fragment to be sequenced, it must have an i7 and i5 primer on either side. Two possible combinations result in non-productive fragments. C) i5 and i7 overhangs are first filled in by a 72°extension step. This filled in sequence is then used as a template for PCR amplification of fragments. D) To begin bridge amplification, PCR products are first denatured into single strands. Next, these single strands are flowed over a lawn of oligos grafted to the flow cell floor which have a complementary sequence that anneals to the P5 or P7 sequences of the index 1 (i7) or index 2 (i5) PCR primers. Once annealed, complementary strands are then polymerized and the original binding strand washed away after being denatured. The single stranded, newly polymerized sequence is then able to bind to a complementary oligo grafted to the flow cell, and the complementary strand is polymerized and subsequently denatured. This process repeats many times to create clonal colonies of each read. Finally, the reverse strands are cleaved and washed away and 3*′* ends of bound reads are blocked. The initial binding and end product stages of bridge amplification are depicted in the panel. E) Sequencing primers are used to determine the sequence downstream of the insertion event. Regardless of orientation, the first base of every read (the end of the adapter sequence) will be 4 positions away from the centrally recognized base.

**Fig. S2.**
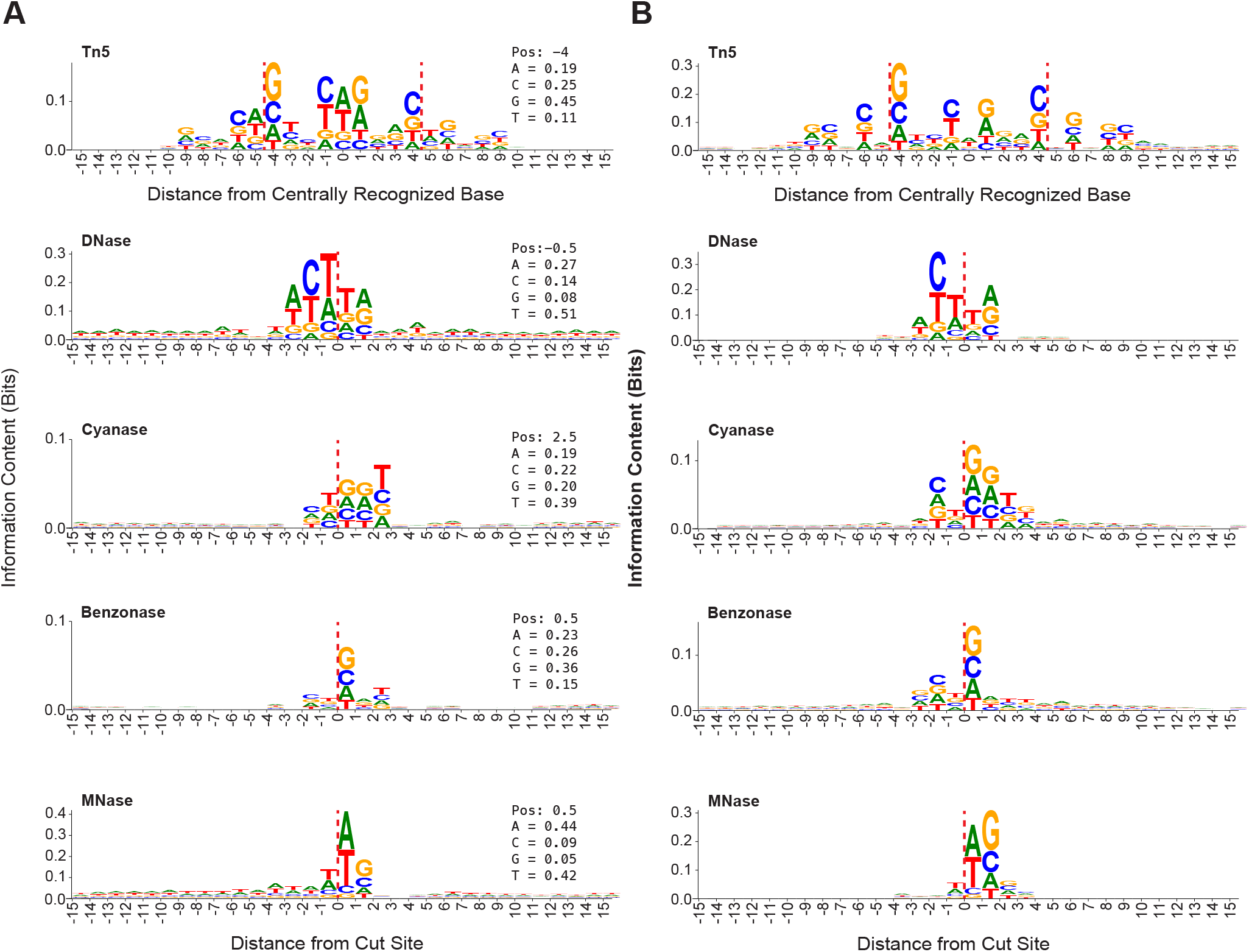
Tn5 sequence bias motif is more complex than nucleases. A) These sequence logos (Gavin E. Crooks and Brenner 2004; Schneider and Stephens 1990) representing enzyme biases were generated with equiprobable background nucleotide frequencies, which does not incorporate information about genomic AT/CG content. Nucleotide frequencies listed in the inset to the right correspond to the highest information content position (Pos). Nuclease cleavage sites are indicated by the dashed red line. B) Plots of enzymatic sequence bias motifs as seen in (Figure 1A), with the y-axis limit scaled to the position with highest information content.

**Fig. S3.**
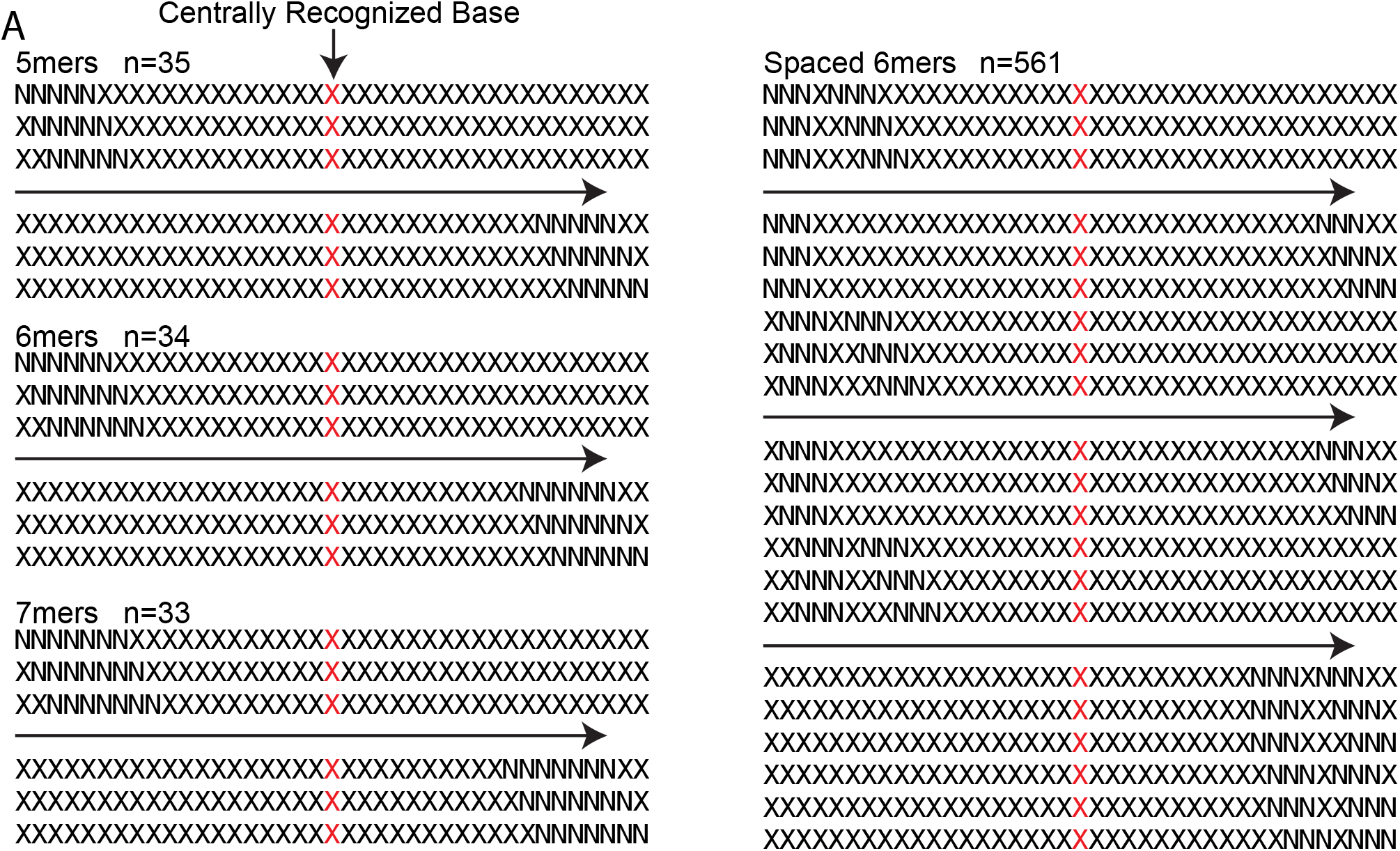
Many 5/6/7-mer and spaced 6-mer combinations were inputs to rule ensemble modeling of Tn5 bias. A) This graphic depicts direct k-mer scaling positions used as input for the Tn5 rule ensemble model. In this example, the positions marked ‘X’ represent unmodeled positions, while the ones marked ‘N’ are the positions which each read is scaled by. The ‘n’ in the upper left corner is the number of permutations per k-mer size. Each graphic for each contiguous k-mer size represents the first three and last three input positions. The graphic for spaced 6-mer positions shows how the leading 3-mer is moved across the span, followed by a single base pair movement of the trailing 3-mer, to sample all possible combinations of spacing between the two.

**Fig. S4.**
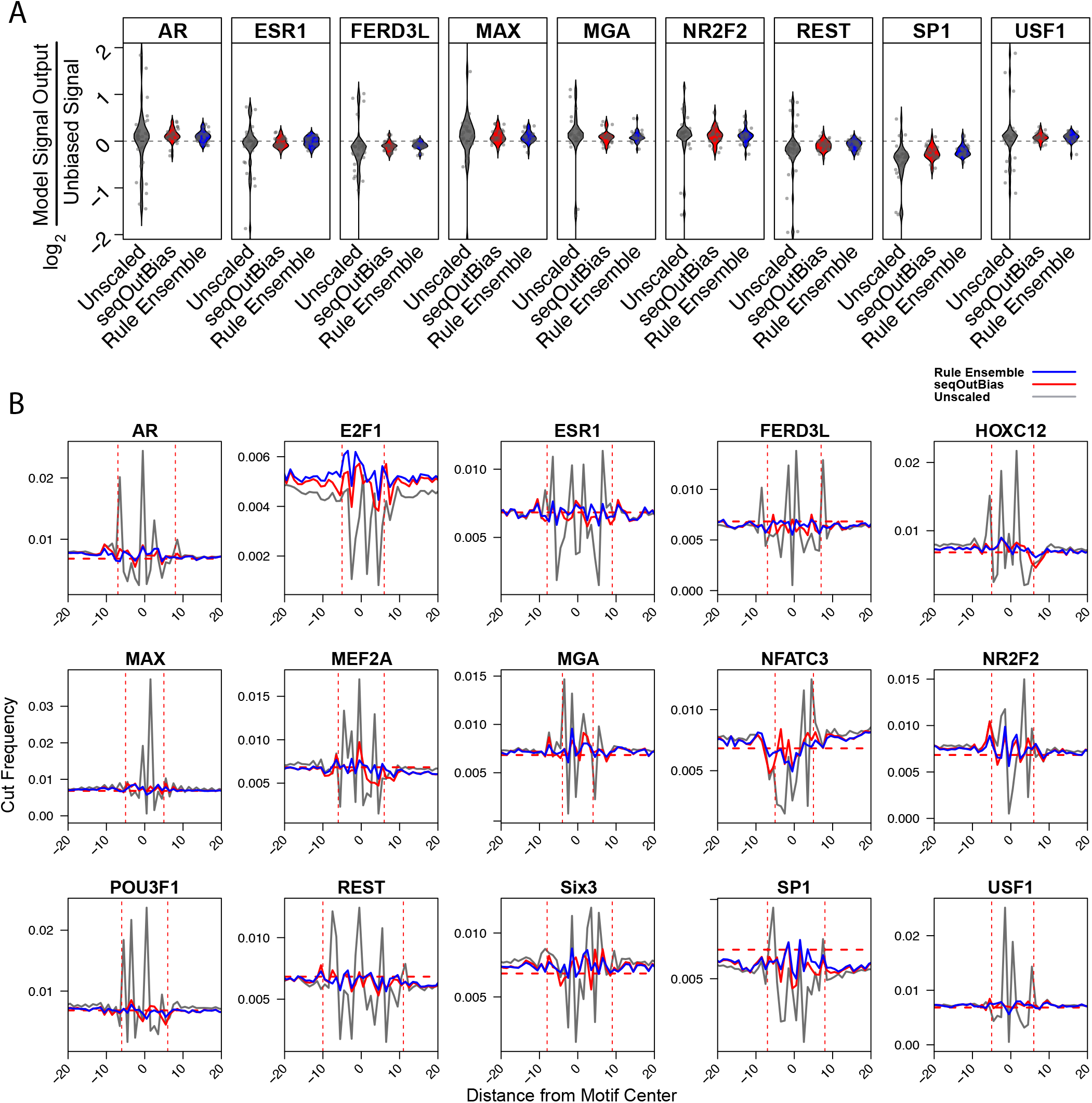
Rule ensemble modeling corrects DNase bias. A) The remaining violin plots which compare the log_2_ of output divided by unbiased signal for each test transcription factor’s composite profile. B) Composite profiles of the test set transcription factors show correction of DNase sequence bias. Each plot shows rule ensemble output compared with direct k-mer scaling of the 5-mer that encompasses the five most influential positions in Figure 5A. The horizontal dashed red lines indicate random cleavage, while the vertical dashed red lines indicate the beginning and end of the motif.

**Fig. S5.**
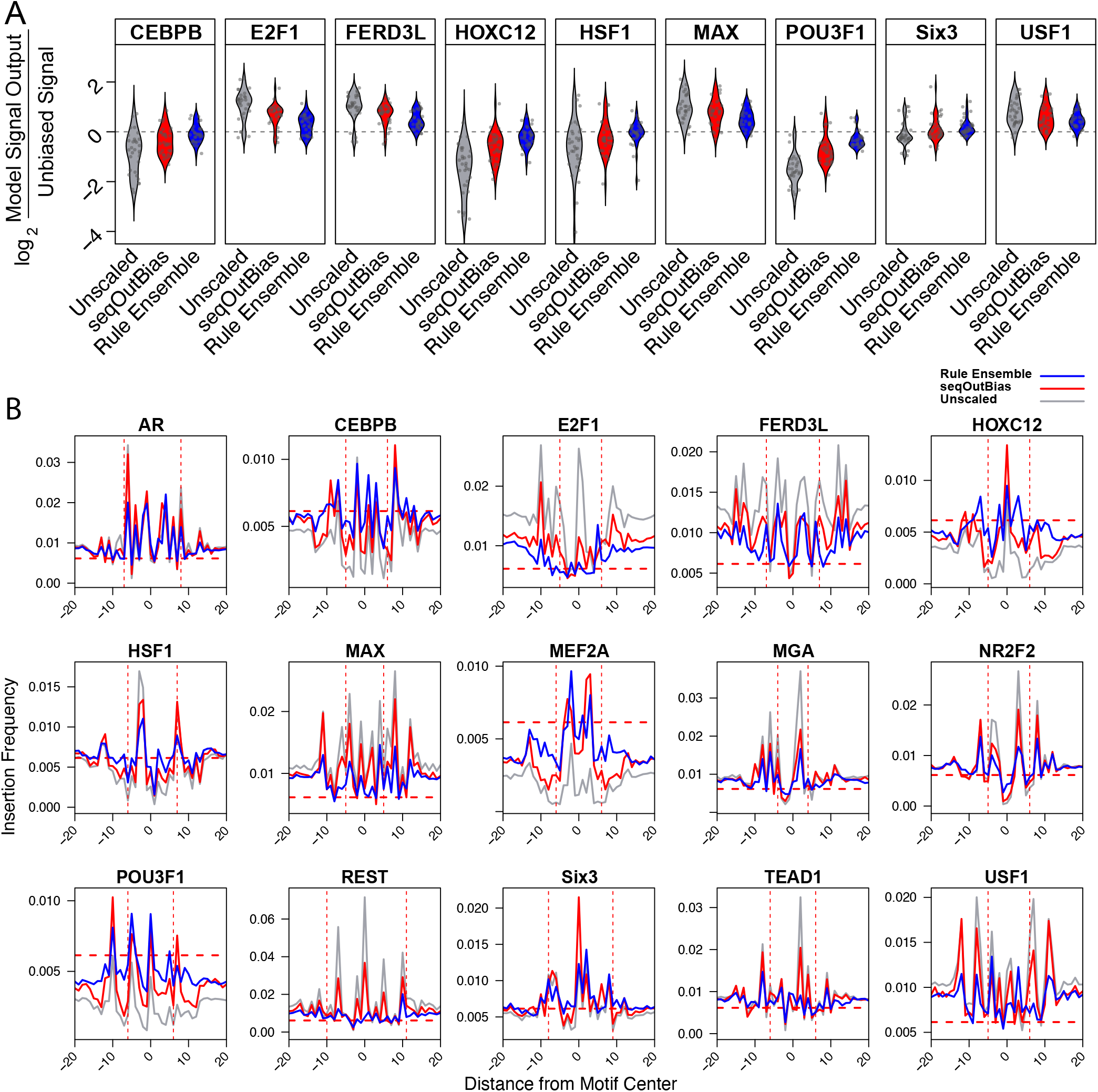
Rule ensemble modeling corrects Tn5 bias. A) The single nucleotide bias correction comparison of the log_2_ ratio of model output to unbiased signal for each transcription factor indicate that rule ensemble outperforms direct k-mer scaling. These motifs represent the remaining test set motifs that are not in Figure 6. B) The composite profiles of transcription factors from rule ensemble output compared with direct k-mer scaling highlight the improvement of rule ensemble modeling. Dashed horizontal red lines denote random cleavage and the dashed vertical lines bookend each transcription factor motif.

